# Genome-wide methylation prediction of biological age using reproducing kernel Hilbert spaces and Bayesian ridge regressions

**DOI:** 10.1101/2020.08.25.266924

**Authors:** Mahmoud Amiri Roudbar, Mehdi Momen, Seyedeh Fatemeh Mousavi, Siavash Salek Ardestani, Fernando Brito Lopes, Daniel Gianola, Hasan Khatib

## Abstract

The use of DNA methylation signatures to predict chronological age and the aging rate is of interest in many fields, including disease prevention and treatment, forensics, and anti-aging medicine. Although a large number of methylation markers have been found to be significantly associated with age, most age-prediction methods use a small number of markers selected based on either previously published studies or datasets containing methylation information. Here, we implemented reproducing kernel Hilbert spaces (RKHS) regression and ridge regression model in a Bayesian framework that utilized phenotypic and methylation profiles simultaneously to predict chronological age. We used over 450,000 CpG sites from the whole blood of a large cohort of 4,409 human individuals with a range of 10-101 years of age. Models were fitted using adjusted and un-adjusted methylation measurements for cell heterogeneity. Non-adjusted methylation scores delivered a significantly higher prediction accuracy than adjusted methylation data, with a correlation between age and predicted age of 0.98 and a root-mean-square error (RMSE) of 3.54 years in non-adjusted data, 0.90 (correlation) and 7.16 (RMSE) years in adjusted data. Reducing the number of predictors through subset selection improved predictive power with a correlation of 0.98 and an RMSE of 2.98 years in the RKHS model. We found distinct global methylation patterns, with significant hypermethylation in CpG islands and hypomethylation in other CpG types including CpG shore, shelf, and open sea (p < 5e-06). Epigenetic drift seemed to be a widespread phenomenon as more than 97% of the age-associated methylation sites had heteroscedasticity. Apparent methylomic aging rate (AMAR) had a sex-specific pattern, with an increase in AMAR in females with age compared to males.

## INTRODUCTION

Aging is a complicated process with time-dependent changes in a set of multiple biological functions and subject to regulation by signaling pathways and transcription factors. Reduced stability of epigenetic patterns over a lifetime in adult somatic tissue is one of the most important progressive deteriorating changes which may increase significant pathologies, including cancer, metabolic disruptions, cardiovascular malfunctions, and neurological disorders, and result in more susceptibility to death in elders (López-Otín *et al*. 2013). This instability is referred to as ‘aging epigenetics’, and associations between age and DNA modifications other than sequence mutations have been investigated (Fraga and Esteller 2007). Recent research has shown a robust relationship between aging and changes in one of the most intensely studied epigenetic modifications, DNA methylation (Heyn *et al*. 2012; Hannum *et al*. 2013; Horvath 2013; Bormann *et al*. 2016; Petkovich *et al*. 2017). DNA methylation changes can impact the aging rate through altering the expression of age-related genes via the silencing of DNA repair mechanisms or silencing anti-inflammatory genes.

Predicting quantitative traits with regression models for dense molecular markers—such as high-throughput genotyping data, sequencing information, and methylation profiling microarrays—is essential in animal and plant breeding schemes (de los Campos *et al*. 2013; Hu *et al*. 2015) and human genetics (Vazquez *et al*. 2016a; Amiri Roudbar *et al*. 2020). When fitting high-density molecular markers in prediction models, the number of predictors dramatically outweighs the number of observations. Several variable selection methods or shrinkage estimation procedures have been proposed for nonparametric or parametric regression models (Gianola *et al*. 2006; de los Campos *et al*. 2013). Most complex traits can be predicted with reasonable accuracy using the infinitesimal model, which assumes a very large number of loci contributing to the trait, each with infinitesimal effect (Buckler *et al*. 2009; Yang *et al*. 2010). Further, a large number of DNA methylation marks show significant associations with age (Heyn *et al*. 2012; Hannum *et al*. 2013), suggesting that the infinitesimal models may also be appropriate for predicting chronological age. Gianola *et al*. (2006) presented a semiparametric procedure, named reproducing kernel Hilbert spaces (RKHS) regression, for the prediction of quantitative traits that makes use of phenotypic and molecular information simultaneously. RKHS has been found to provide a robust framework for integrating high-dimensional multilayer omic-data for the prediction of survival after diagnosis of breast cancer (Vazquez *et al*. 2016b). Bayesian methods are powerful alternative approaches for predicting complex traits using variable selection and shrinkage of estimates. Various Bayesian parametric linear regressions that differ in their priors have been proposed for genome-enabled prediction (see Gianola (2013) and de los Campos *et al*. (2013) for discussions of Bayesian regressions). Bayesian ridge regression (BRR) performs a homogenous shrinkage across markers. Statistically, BRR on markers is equivalent to genomic best linear unbiased prediction (GBLUP), but with variance parameters estimated Bayesianly (Vanraden 2008; Gianola 2013).

The use of DNA methylation signatures to predict chronological age and aging rate is of interest to many fields, including disease prevention and treatment, forensics, and evaluation of anti-aging medicine. In recent years, many methods have been proposed to predict an individual’s chronological age (Bocklandt *et al*. 2011; Hannum *et al*. 2013; Bekaert *et al*. 2015; Naue *et al*. 2017; Vidaki *et al*. 2017). However, most of these studies used a small number of markers selected based on previously published studies or used methylation information retrieved from 27K or 450K microarrays for marker selection. Selected markers, ranging from 3 to 513, have been used for age prediction using as multivariate linear regression models (Hannum *et al*. 2013; Hong *et al*. 2017; Levine *et al*. 2018) or nonparametric methods such as machine learning algorithms (Xu *et al*. 2015; Vidaki *et al*. 2017).

The power of semiparametric and Bayesian approaches for age prediction has not yet been evaluated. Here, we evaluated RKHS and BRR models applied to whole methylome prediction of age in humans using a large cohort of 4,409 individuals composed of four publicly available DNA methylation datasets representing a range of 10-101 years of age. Furthermore, marker subset selection was carried out using epigenome-wide association study (EWAS) results to retrieve age-associated methylation sites, and Breusch-Pagan test (BPtest) results were used to retrieve age-associated methylation sites without heteroscedasticity. Finally, to better understand how the methylome changes with age, we evaluated genome-wide methylomic profiling with this large cohort of individuals, including the effect of sex.

## MATERIAL AND METHODS

### Datasets

Methylation data were obtained from four different datasets. All data are available in Gene Expression Omnibus (GEO) with accession numbers “GSE55763”, “GSE40279”, “GSE50660” and “GSE56105” (see Supplementary file 1 Table S1 and Figure S1). DNA extraction, preparation procedures, and DNA methylation profiling were previously described for each dataset separately(Hannum *et al*. 2013; Liu *et al*. 2013; McRae *et al*. 2014; Tsaprouni *et al*. 2014; Lehne *et al*. 2015). Briefly, to assess the methylation levels of over 485,000 CpG sites per sample, DNA extracted from whole blood was treated with bisulfite and then hybridized to the Illumina Infinium 450k Human Methylation Beadchip. In each dataset, individual measured methylation values with detection p-value > 0.01 were set as missing values. Probes unsuccessfully measured in 5% of samples, with SNPs at CpG or single-base extension (SBE) sites and located on the X and Y chromosomes were excluded from the analyses. Cross-reactive probes were also removed. Missing values were imputed using the R “impute” package with ten nearest neighbors averaging (Troyanskaya *et al*. 2001). After data preprocessing, all datasets were merged, and probes that were common in all datasets were kept. In the end, 423,394 methylation sites from 4,409 individuals remained for training and validation of the models. For each CpG site, the beta value was used to indicate the percentage of methylation. The values ranged from zero to one for completely unmethylated and fully methylated, respectively. After merging the datasets, we used the z-score conversion to normalize the methylation levels.

### Estimating cell heterogeneity for each sample

We used the Houseman *et al*. (2012) regression calibration approach algorithm to estimate the relative proportion of pure cell types. For each sample, the proportions of white blood cell types (WBC), including granulocytes, monocytes, B cells, CD4+ T cells, CD8+ T cells, and natural killer cells, were inferred using 473 CpGs which had previously shown cell-type-specific methylation patterns (Reinius *et al*. 2012). This was performed using the R function projectWBC.

### Statistical models for predicting chronological aging

Parametric and semiparametric approaches were used to predict chronological aging. A statistical and computational challenge was that the number of methylation sites exceeds the number of data points. Therefore, the shrinkage estimation in prediction models was applied (Meuwissen *et al*. 2001; Gianola *et al*. 2006). The predictive ability of either the entire methylation data set and of subsets was evaluated using RKHS regression (Gianola and van Kaam 2008; Morota and Gianola 2014), and Ridge Regression (Hoerl and Kennard 1970) (BRR) approaches based on a Bayesian framework.

### Reproducing kernel Hilbert space (RKHS)

RKHS regression is a powerful semiparametric approach to cope with the issues of dimensionality and complexity raised by a considerable number of predictors (de los Campos *et al*. 2010). In RKHS, we built a covariance structure among methylation values. We treated age as a continuous response in the following kernel regression model:

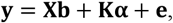

where **b** is a vector of fixed effects including an intercept and sex and dataset effects with associated incidence matrix **X, K** is an *n* × *n* positive definitive kernel matrix indexed by adjusted or non-adjusted methylation levels, **α** is the vector of RKHS regression coefficients estimated as the solution of 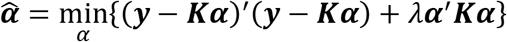, where *λ* is a regularization parameter, and **e** is the vector of residual effects. Here ***α*** and **e** were assumed as random vectors with distributions 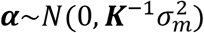 and 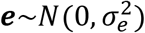, where 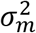, and 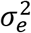 are the methylomic and the residual variances, respectively. RKHS was fitted using Gaussian kernels. The Gaussian kernel function was:

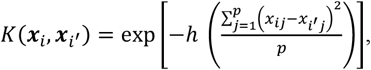

where 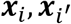 are vectors of methylation measurements in the *i* and *i*’ individuals (*i* = 1, 2,…, n) for *p* number of methylation sites (*j* = 1,…, *p*), *h* is a bandwidth parameter chosen with cross-validation, based on their ability to predict testing set.

### Bayesian Ridge Regression (BRR)

To assess the parametric method, we used a whole-genome regression (Meuwissen *et al*. 2001) approach based on a Bayesian-regression framework. In this model, a Gaussian prior density, or BRR, was assigned to methylation effects to control the shrinkage of estimates. We considered age as a continuous response with the following model equation in matrix notation:

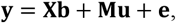

where **M** is a matrix of the non-adjusted beta value in *i*th individual at *j*th methylation site (*j* = 1,…, p), **u** is a vector for the corresponding methylation site effect. With assuming the mentioned model and assumptions, the conditional distribution of data is as follow:

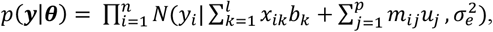

where ***θ*** is the vector of unknown parameters including the intercept, regression coefficients, and residual variance with the following prior density:

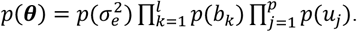

Here, the fixed effects are assigned as a flat prior, the residual variance is assigned as a scaled-inverse χ^2^ density with degrees of freedom df_e_ and scale parameter S_e_ using the default treatment of variances implemented in the BGLR R package^39^, and methylation effects assigned as a Gaussian density.

### EWAS subset selection

In the full models used, there were 4,409 samples with more than 420,000 methylation sites as predictor variables. Fitting this large number of methylation covariates is computationally expensive. A previous association test study involving methylation sites and chronological age found up to 15% significant methylation sites(Hannum *et al*. 2013). Here, we applied an EWAS subset selection approach to assess the impact of the number of predictors in the prediction model. Accordingly, we fitted the age-associated methylation positions (aDMPs) only and also evaluated randomly selected methylation sites. For each cross-validation (CV) sample, we conducted an EWAS to find aDMPs and contrasted the predictions with those attained with random subsets of methylation probes.

### BPtest subset selection

Each aDMP was tested for heteroscedasticity using the BPtest model (Breusch and Pagan 1979). Methylation trends due to age, sex, and dataset were removed by fitting a linear model for each aDMP, and the residuals were retrieved. Next, the BPtest was performed by fitting a linear regression model to these residuals. aDMPs without heteroscedastic disturbances were selected based on a Benjamini-Hochberg’s false discovery rate (FDR) less than 0.05 (Benjamini and Hochberg 1995). The influence of this selection approach on prediction accuracy was evaluated against randomly selected probes in each CV.

### Subset selection of CpG Sites based on previous studies

We used results from the four studies to fit prediction models based on their candidate CpG sites (Hannum *et al*. 2013; Horvath 2013; Horvath *et al*. 2018; Levine *et al*. 2018). The “optimal” model from these studies selected 71 (Hannum method) (Hannum *et al*. 2013), 353 (Horvath multi-tissue method) (Horvath 2013), 391 (Horvath skin method) (Horvath *et al*. 2018), and 513 (Levine method) (Levine *et al*. 2018) methylation markers that were highly predictive of age. We missed some of these methylation sites due to our quality control. The number of missed CpG sites and their designations are shown in Supplementary File 1, Table S2. Measurements of these candidate CpG sites in each method were used to predict chronological age using the multivariate regression model:

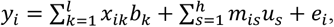

Where *u_s_* is the methylation effect for the sth site from the subset selected based on the four methods listed above.

### Prediction accuracy

We used two measures of predictive accuracy. The first measure was the Pearson correlation coefficient between predicted age and chronological age. The second measure was the root mean square error (RMSE), which is in the same units as the response variable. RMSE measures closeness between chronological and predicted ages and correlation measures association.

To assess the prediction accuracy of chronological age in each model, 5-fold crossvalidations (CVs) were performed. The CV was repeated four times, producing a total of 20 samples. In this case, the entire data was partitioned into five sets. For each CV, 5000 posterior samples were used to compute posterior means of the parameters. The averages of correlation and RMSE between real age and predicted age from the 20 CVs were used to compare the models. All prediction models were fitted using the “BGLR” R package (Pérez and de los Campos 2014). We used the coda package diagnostic tests of convergence in our Bayesian analyses (Plummer *et al*. 2006).

### The identification of age-associated differentially methylated positions (aDMPs)

To identify age-associated methylation positions, we fitted a linear regression of chronological age on each CpG site separately. Sex and dataset were treated as fixed effects in the model. As methylation levels may be affected by the heterogeneity of cell populations, the effects of differential cell count on aDMPs were examined using adjusted and non-adjusted beta values. To control false positives at the potential expense of more false negatives, a stringent Bonferroni-adjusted threshold was used to correct for multiple testing (0.05/423,394 = 1.18e-7).

### Mapping CpG sites and dynamic changes through the aging process

We used IlluminaHumanMethylation450kanno.ilmn12.hg19 package from Bioconductor (Gentleman *et al*. 2004) to group the CpG probes located on the same island, shore, shelf, or open sea. Methylation markers in each CpG type were classified into two groups: 1) the age-associated and 2) not age-associated methylation sites. To evaluate the differences in DNA methylation patterns between the groups with aging, we fitted a linear regression of the average of individual methylation levels in each group at different locations in the genome.

### Functional annotation analyses

Functional annotation for the selected methylation sites was performed using Pathway and disease association analyses. For each methylation site, the R package minfi was used to access annotation for each position (Aryee *et al*. 2014). Then, genes associated with aDMPs were characterized employing BioMart web services through the R package biomaRrt (Durinck *et al*. 2009). Gene lists retrieved from each group were uploaded to the Database of Annotation, Visualization, and Integrated Discovery (DAVID; david.abcc.ncifcrf.gov) v6.8^42^ to link them to associated diseases and Pathway using genetic association database (GAD) source (Becker *et al*. 2004) and KEGG pathway maps (Kanehisa and Goto 2000), respectively.

### The identification methylation sites influencing sex-specific apparent methylomic aging rate (AMAR)

Apparent methylomic aging rate (AMAR) for each individual was computed using results of the EWAS subset selection model, but without a sex variable. The AMAR was calculated as the individual’s predicted age, divided by the actual age. It has been previously shown that sex can affect the aging rate derived from a methylome profile, with a faster aging rate in males than in females (Hannum *et al*. 2013). Methylomic aging rate in males was 4% faster than in females ^3^. Here, we included sex as a dummy explanatory variable to account for a sex effect on methylation levels across chronological ages. To test differences between male and female methylome aging rates, first, we fitted three linear models as follows:

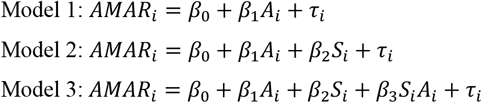

Where *AMAR_i_* is an estimated dependent variable for the *i*th individual; *A_i_* is chronological age for the *i*th individual; *S_i_* is the dummy explanatory variable for the sex; *β*_0_, *β*_1_, *β*_2_, and *β*_3_ are the intercept, age, male effect, and interactions between sex and age (difference between male and female slope over aging), respectively; and *τ_i_* is an independent normal model residual with mean zero and variance 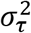. Models were compared using Akaike information criterion (AIC) (Akaike 1998). Model 1 states that there is no difference between males and females in AMAR over aging, but Models 2 and 3 state that there are differences as well as interactions between sexes over aging, respectively.

Second, differences in methylome changes over a wide age range in males and females were tested at each aDMP. Slopes were compared using the model:

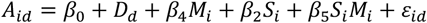

Where *A_id_* is the age for the *i*th individual in the *d*th dataset, *D_d_* is the fixed effect of the *d*th dataset, *M_i_*, is methylation level for the *i*th individual, *S_i_M_i_* is a joint-effect covariate. *β*_4_ and *β*_5_ are methylation effect and interactions between sex and methylation levels over aging, respectively, and *ε_ij_* is an independent normal residual with mean zero and variance 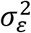. Sex-specific aDMP associations were corrected for multiple testing using the inclusion threshold based on the Benjamini-Hochberg with the desired FDR less than 0.05 (Benjamini and Hochberg 1995).

## RESULTS AND DISCUSSION

### Correcting for cellular heterogeneity

Methylome profiles of 423,394 CpG sites in 4,409 individuals from four different datasets with age varying from 10 to 101 years were obtained. Each methylation measurement was converted to beta values ranging between zero and one for completely-unmethylated and methylated sites, respectively. The methylation profiles were measured in extractions from whole blood samples with heterogeneous cell proportions, which can act as a potential confounder when investigating DNA methylation differences over a wide age range (Jaffe and Irizarry 2014). As shown in previous studies, adjustment for cell proportions can reduce association signals; therefore, it is essential to consider cell composition variability in epigenetic studies based on whole blood and other heterogeneous tissue sources (Liu *et al*. 2013; Jaffe and Irizarry 2014). We tested the association of each cell type proportion with age using a linear model with sex and dataset considered as fixed effects. Cell type proportion was not associated with aging (p > 0.05, Supplementary file 1 table S3-S8); therefore, it was not included in the final model for EWAS, which employed the non-adjusted methylation data.

### Whole methylome prediction model

Epigenetic drift represents the tendency for increasing discordance between epigenomes over aging, and the epigenetic clock refers to specific sites that are consistently related to age (Jones *et al*. 2015). Recent studies have used elastic net regression models and artificial neural network modeling predicting chronological age using methylation clock sites (Hannum *et al*. 2013; Horvath 2013; Petkovich *et al*. 2017; Stubbs *et al*. 2017; Vidaki *et al*. 2017). These methods automatically select a small subset of methylation markers (maximum a few hundred) to fit a model that is highly predictive of chronological age. In contrast, our study found that the number of methylation sites related to age is over 100,000. Then, it appears more sensible to use an age prediction model where all methylation sites are fitted simultaneously. Consequently, we compared two parametric and semiparametric methods. Bayesian ridge regression (BRR) was used as a parametric method to fit a huge number of methylation sites by assuming a Gaussian prior distribution of the effect of each methylation site (de los Campos *et al*. 2013). We chose reproducing kernel Hilbert spaces regression (RKHS) as a semiparametric method, with a Gaussian kernel used for the prediction of chronological age (Morota and Gianola 2014). Methylomic age could be sensitive to many factors such as sex, which can significantly influence methylomic aging rate (Hannum *et al*. 2013). Accordingly, we first performed epigenome-wide prediction using adjusted and non-adjusted beta values according to the proportion of white blood cells, sex, and dataset using a linear model. Then, different subsets of methylation markers, including EWAS, BPtest, and randomly selected markers were investigated.

### Effect of adjustment for estimated white blood cell heterogeneity on age prediction accuracy

As whole blood is a heterogeneous ensemble of white cells, with each type having its own epigenetic profile, adjustment for white blood cell heterogeneity in EWA studies is strongly recommended when this source of tissue is used. Heterogeneous cell populations in the blood may act as a potential confounder when cell distribution differs over target traits. For instance, adjustment for white blood cell proportions reduced the strength of association signals in rheumatoid arthritis disease (Liu *et al*. 2013). Although EWASs have illustrated the importance of adjusting for changes in cell-type composition (Liu *et al*. 2013; Jaffe and Irizarry 2014), the impact on whole methylation prediction hasn’t been studied in age prediction. However, adjusting methylation profiles for cellular heterogeneity had a detrimental effect on the classification ability of rheumatoid arthritis cases from controls (Amiri Roudbar *et al*. 2020). Here, we conducted whole methylome prediction analyses using methylation measurements adjusted and non-adjusted for cell heterogeneity to evaluate the effect on prediction accuracy of chronological age using the RKHS approach. The use of non-adjusted methylation produced higher prediction accuracy than adjusted methylation data (Table 1). For this reason, we used non-adjusted methylation data for subsequent analyses.

**Table 1.**
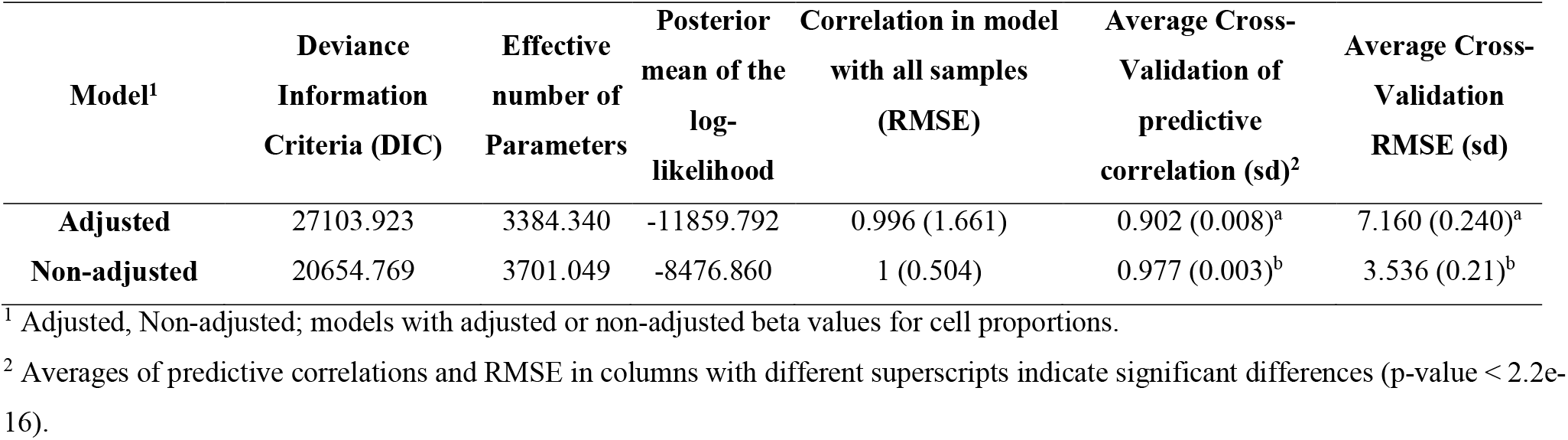
Goodness of fit, prediction accuracy, and root mean square error (RMSE) of the reproducing kernel Hilbert space (RKHS) model including all methylation sites.

### Predictive ability of different methylation subsets

Model complexity, prediction accuracy, and goodness of fit for each type of subset selection are given in Table 2. Using EWAS, we found that most methylation sites showed no association with age. Although BRR and RKHS use shrinkage of estimates to handle the problem of a large number of the predictors relative to sample size, subset selection was used to assess the impact of the pre-selection of predictors and then reduce the number of predictors. EWAS subset selection seemed to improve prediction accuracy, as the lowest RMSE and the highest correlation were achieved in this subset. Using EWAS selection, there were no significant differences between semiparametric and parametric approaches. The correlation of predicted age between the two approaches was very high (Figure S2). These results are in agreement with a previous study that found no difference in maize in whole-genome prediction accuracies between parametric and semiparametric approaches (Riedelsheimer *et al*. 2012). Comparing EWAS subset selection with random selection, the CV analyses indicated the usefulness of EWAS subset selection. Our accuracy was a little higher than reported in previous studies that used the elastic net approach (Hannum *et al*. 2013; Horvath 2013). This increase in accuracy could be due to the large sample size used in our study, fitting all aDMPs simultaneously, or the different methodologies used in these studies.

**Table 2.** Goodness of fit, prediction accuracy, and root mean square error (RMSE) reproducing kernel Hilbert space (RKHS) and Bayes ridge regression (BRR) models fitted to subsets of the methylation sites, non-adjusted beta values.

| Model<sup>1</sup> | Deviance<br>Information<br>Criteria (DIC) | Effective<br>number of<br>Parameters | Posterior mean<br>of the log-<br>likelihood | Correlation in<br>model with all<br>samples (RMSE) | Average Cross-Validation of<br>predictive<br>correlation (sd)<sup>2</sup> | Average<br>Cross-Validation<br>RMSE (sd) |
| --- | --- | --- | --- | --- | --- | --- |
| **EWAS Subset RKHS** | 19720.036 | 3474.908 | -8122.564 | 0.999 (0.566) | 0.984 (0.002)<sup>a</sup> | 2.983 (0.155)<sup>a</sup> |
| **EWAS Subset BRR** | 19717.638 | 3494.274 | -8111.682 | 0.999 (0.554) | 0.983 (0.002)<sup>a</sup> | 2.999 (0.183)<sup>a</sup> |
| **Random EWAS Subset RKHS** | 20689.105 | 3687.112 | -8500.997 | 1 (0.518) | 0.977 (0.003)<sup>b</sup> | 3.588 (0.210)<sup>b</sup> |
| **Random EWAS Subset BRR** | 20744.651 | 3680.213 | -8532.219 | 1 (0.529) | 0.976 (0.003)<sup>b</sup> | 3.592 (0.234)<sup>b</sup> |
| **BPtest Subset RKHS** | 24776.629 | 2525.258 | -11125.686 | 0.993 (1.940) | 0.961 (0.003)<sup>c</sup> | 4.550 (0.143)<sup>c</sup> |
| **BPtest Subset BRR** | 26008.048 | 1537.348 | -12235.350 | 0.982 (3.121) | 0.959 (0.003)<sup>d</sup> | 4.710 (0.157)<sup>c</sup> |
| **Random BPtest Subset RKHS** | 23938.746 | 3155.407 | -10391.670 | 0.997 (1.302) | 0.955 (0.004)<sup>e</sup> | 4.908 (0.189)<sup>d</sup> |
| **Random BPtest Subset BRR** | 25814.363 | 1674.056 | -12070.154 | 0.984 (2.931) | 0.952 (0.004)<sup>f</sup> | 5.068 (0.238)<sup>d</sup> |

<sup>1</sup> Random EWAS/BPtest Subset; models with randomly chosen in number of EWAS/BPtest subset.
<sup>2</sup> Averages of predictive correlations and RMSE with different letters indicate significant differences (p-value < 0.01).

It has been suggested that individuals have different methylomic aging rates (Hannum *et al*. 2013) and that DNA methylation changes accumulate over the lifetime. Hannum *et al*. (2013) reported 27,800 markers with heteroscedasticity, of which 99.8% showing an increase in variation with age. We tested each aDMP for heteroscedasticity using a BPtest, and excluded aDMPs with non-normally distributed and serially independent residuals. Markers whose residual variance showed a change with age are illustrated in Figure 1. Of 123,930 aDMPs, 121,023 markers showed heteroscedasticity (FDR < 0.05), and more than 97% of them showed an increase for independent residuals with age (Supplementary file 1, Table S9). These results indicate that epigenetic drift is a more extensive phenomenon than generally believed. For BPtest subset prediction, after excluding aDMPs with heteroscedasticity, the estimate of prediction accuracy from CV analyses was relatively high. Average correlations between age and predicted age retrieved from 20 CVs were 0.96 (with 4.55 years RMSE) and 0.96 (with 4.71 years RMSE) for RKHS and BRR, respectively. These estimates were about 0.7% higher than those when the same number of randomly selected methylation sites (random BPtest subset) was used. In this subset selection method, prediction accuracies were 0.2% (significantly) higher for RKHS, than for BRR was used, but did not differ in RMSE. Selecting subsets of methylation markers with EWAS produce a gain in prediction accuracy.

**Figure 1.**
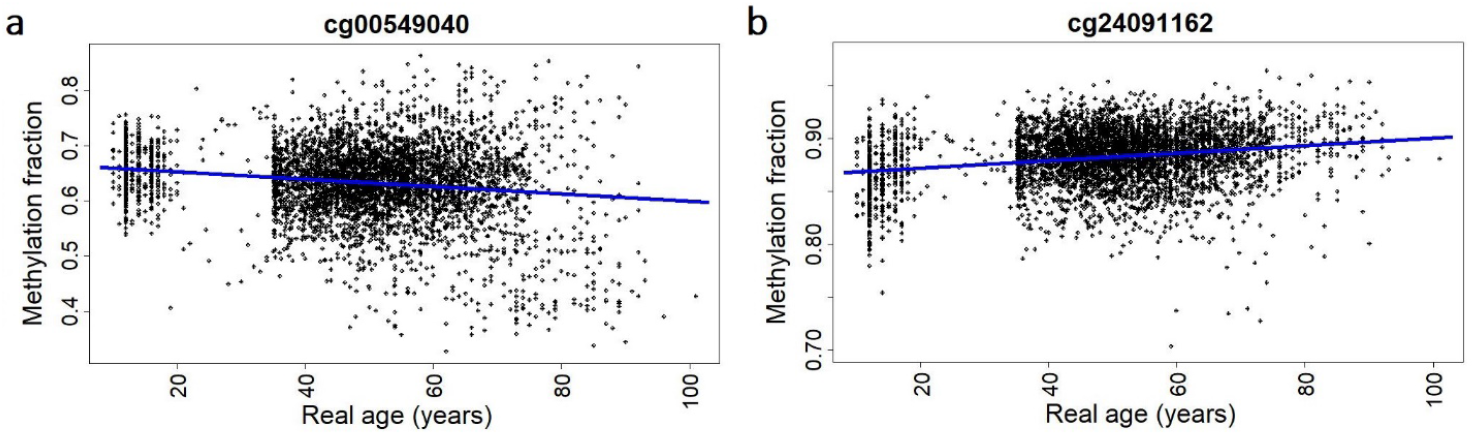
Plots of Methylation beta values for the markers with (a) and without (b) heteroscedasticity. Plots show changes in methylation residuals with age, defined as the difference between an individual’s methylation beta value and expected methylation beta value (blue line).

### Predictive ability of methylation site subsets based on previous studies

To further evaluate whole methylome prediction using RKHS and BRR, four previously presented procedures—Hannum method, Horvath multi-tissue, Horvath skin method, and Levine method— were also evaluated. We applied these methods using adjusted and non-adjusted methylation measurements for cell heterogeneity. The correlation between chronological age and predicted age using adjusted (non-adjusted) measures were 0.57 (0.55), 0.57 (0.55), 0.56 (0.54) and 0.57 (0.55) for Hannum, Horvath multi-tissue, Horvath skin and Levine methods, respectively (Figure 2a). RMSE was 14.9, 14.9, 15.2, and 15.0 years, respectively, when the adjusted methylation measurement was used (Figure 2b). These accuracies were significantly lower than those obtained using RKHS and BRR (P < 0.05). Fitting all methylation sites simultaneously yielded better predictions of age.

**Figure 2.**
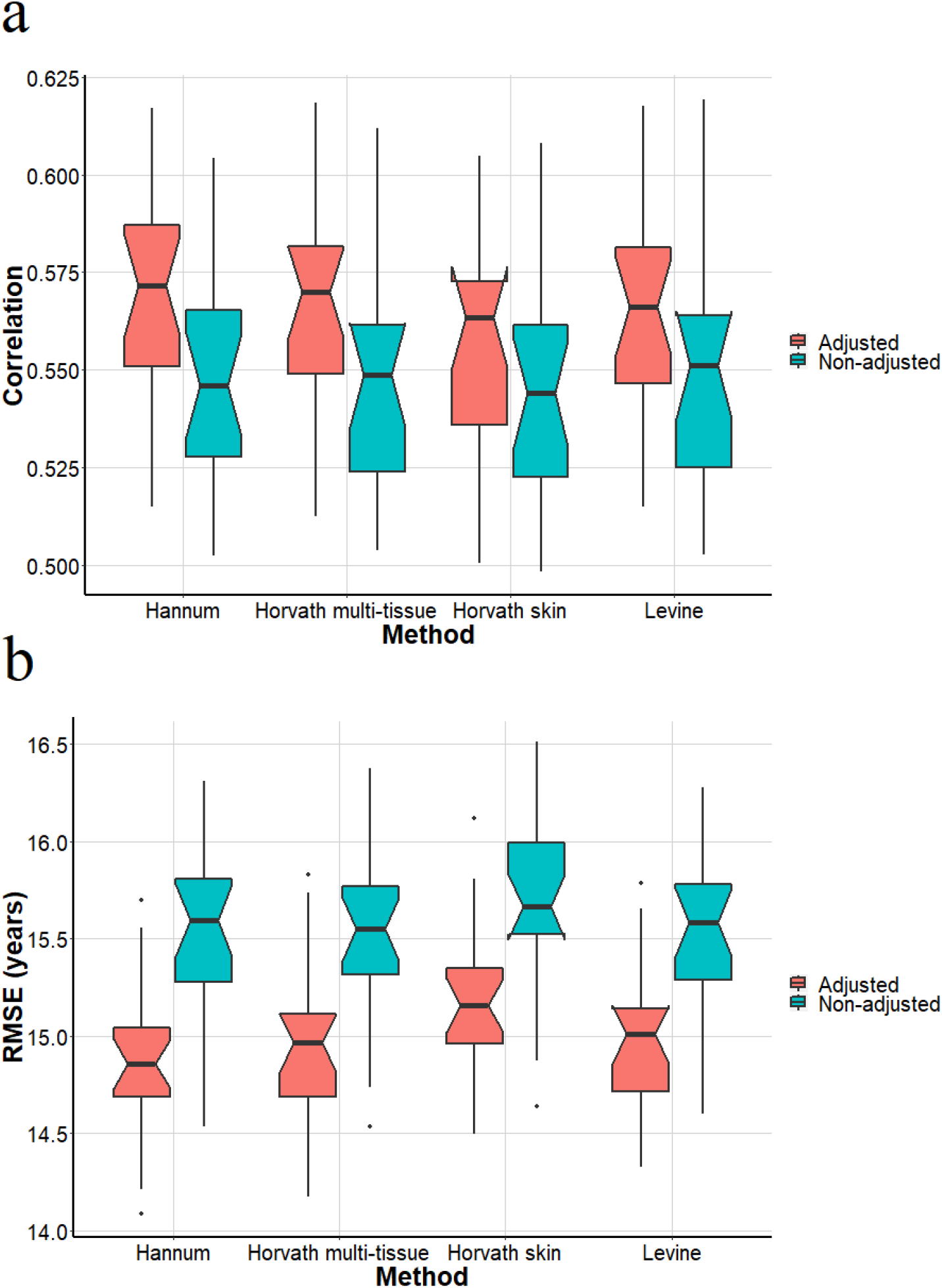
Boxplot of average cross-validation of predictive accuracy (a) of correlation and root mean square error (RMSE) (b) between chronological and predicted age for four methods using adjusted and non-adjusted methylation measurements.

### Age-associated methylation signatures

After quality control on methylation profiles, two association tests were evaluated for the cell proportion correction effects. Results indicated that the number of probes, significantly associated with age, was reduced from 123,930 probes in non-adjusted data to 107,806 probes in adjusted EWAS (Figure 3). The number of aDMPs represents the largest number of methylation sites that can be used for the study of aging. These huge numbers of age-associated markers point to the complexity of aging and can be used for future studies.

**Figure 3.**
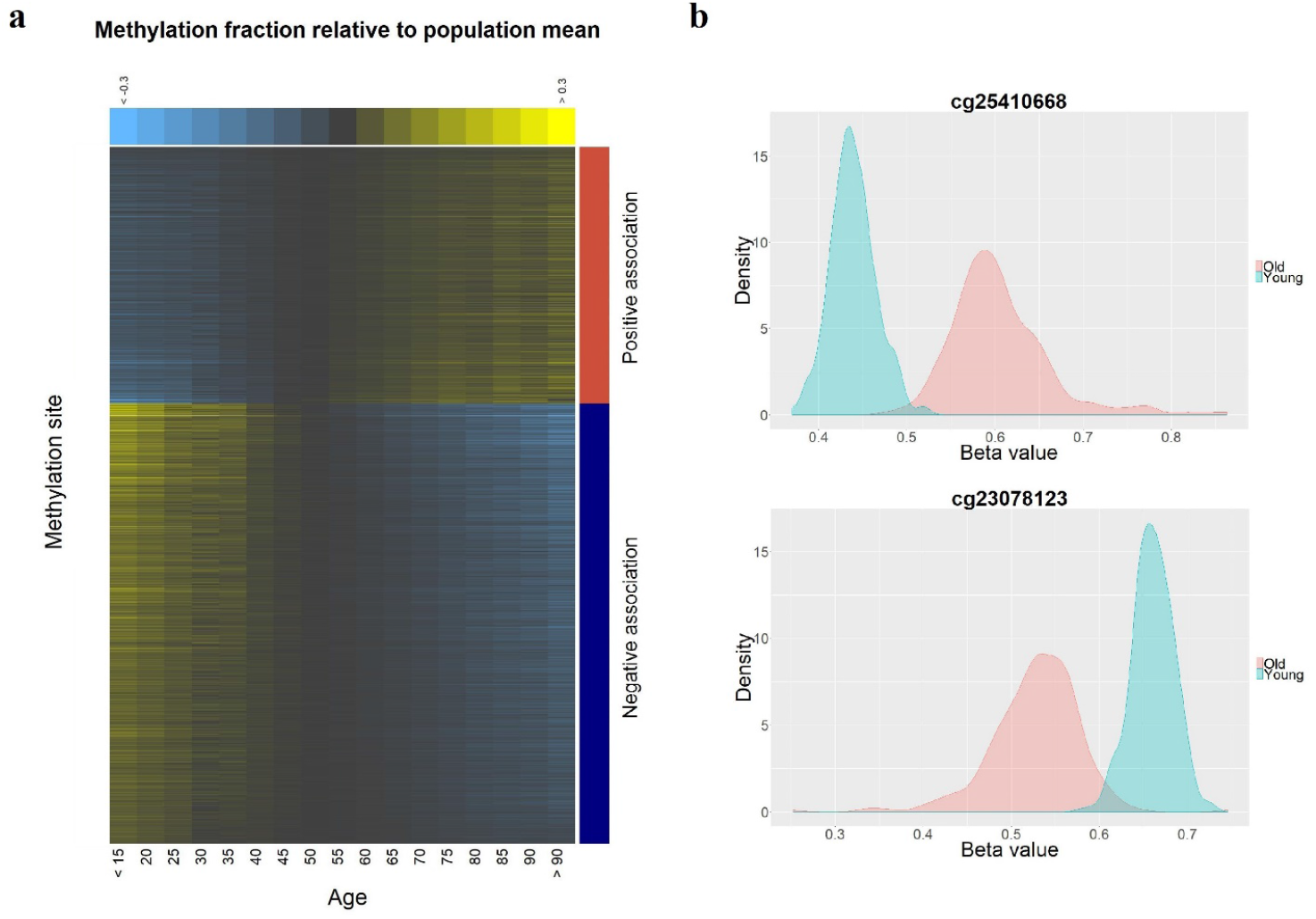
Non-adjusted methylation profiles by age. a) Heatmap of the top 1000 age-associated methylation markers. CpG sites are sorted by the magnitude of regression coefficients. Individuals are grouped into intervals of 5 years and ordered from the youngest to the oldest group. b) Density plots of beta values by age group for positively (top) and negatively (bottom) age-associated methylation markers in the 10% youngest (iris blue) and 10% oldest (salmon) individuals.

### Dynamic changes of the methylome with aging

CpG islands (CGIs) are, on average, 1,000 base pairs (bp) long and can be distinguished from other genomic regions by being GC-rich, CpG-rich and mostly unmethylated. They are frequently associated with more than 70% of the promoter region of genes (Deaton and Bird 2011). CpG shores are considered as a sequence up to 2 kb distant from CGIs. It has been shown that the most tissue-specific differential methylation in normal tissues occurs more frequently in CpG shores than in CpG islands (Irizarry *et al*. 2009). The CpG regions were further classified by including CpG shelves as sequences 2 to 4 kb distant from CGIs and CpG open sea as more than 4 kb distant from CGIs (Bibikova *et al*. 2011). Global DNA methylation level decreases with aging in mammalian tissues (Wilson and Jones 1983). It has been reported that the methylation of a CGI located in the promoter region of the estrogen receptor (ER) gene was associated with age in normal colorectal mucosa (Issa *et al*. 1994). Further, a vast proportion of age-related methylation sites in normal breast tissue was located in CGIs, and a close relationship between age-related DNA methylation changes and epigenetic alterations was present in breast tumors (Johnson *et al*. 2014). In a study on a cancer-free population, no significant differences in global DNA methylation were found between different middle age groups (Zhang *et al*. 2011). In this study, EWAS results revealed that about 30-33 % of age-associated methylation sites were found in adjusted and non-adjusted data and were located in CGIs with a higher proportion of positive association (83%, see Supplementary file 1, Table S10). The proportion of positive associations drastically decreased in CpG shore, shelf, and open sea areas in the adjusted and non-adjusted beta values. These results indicate that methylation sites located in CGIs tend to be hypermethylated with increasing age, whereas aDMPs located in other CpG types, including shore, shelf, and open sea areas, tend to be hypomethylated with aging. To investigate methylation patterns in different types of CpG sites, we calculated the average of age-associated and not-associated methylation beta values for each CGI type separately and evaluated changes of average methylation with aging (Figure 4).

**Figure 4.**
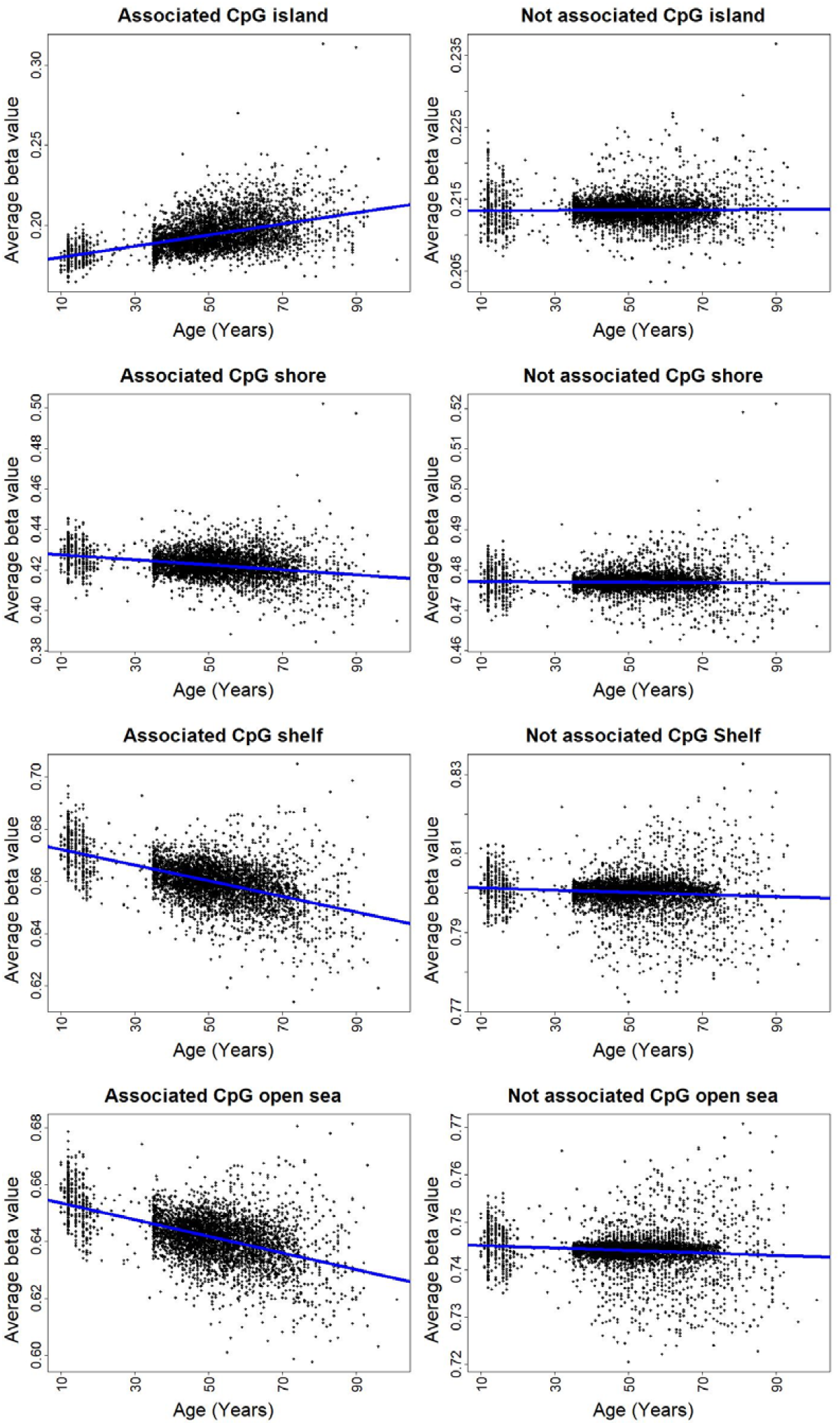
Plots of regressions average of individual methylation levels at different locations in the genome on age. The plots illustrate the association of average beta values and age calculated from the four CpG types, including CpG island (a), shore (b), shelf (c), and open sea (d). Methylation markers in each CpG type classified into two categories: age-associated and not-associated methylation sites. In age-associated sites, the global methylation level increased and decreased with age in CpG island and Off CpG island, respectively (p < 5e-06). In not-associated methylation sites, CpG shelf and open sea were significantly demethylated with increasing age (p < 0.01).

Interestingly, the average methylation level of aDMPs located in CpG islands was significantly higher in older individuals. With increasing distance from CpG island, the average methylation levels for aDMPs significantly decreased with aging. These results are in agreement with previous studies that found age-related methylation sites to be hypermethylated and hypomethylated preferentially at CpG islands (Christensen *et al*. 2009) and CpG shores (Horvath 2013), respectively. The average methylation level for non-associated methylation sites changed slightly and remained constant over the age range. These results enabled us to identify distinct age methylation patterns in CpG types according to distance from CGIs. This global methylation pattern revealed that age-related changes in DNA methylation are not distributed randomly on the genome. Understanding the mechanisms that produce these location-specific methylation patterns during the human lifespan warrant further research.

Recently, a significant difference in the average methylation levels between males and females was observed in individuals ranging 41 to 55 years of age (Tsang *et al*. 2016). A significantly lower methylation level was found for males than for females. In contrast, a study of human peripheral blood from persons ranging from 45 to 75 years, females had lower methylation levels (Zhang *et al*. 2011). With these contradictory reports, the global DNA methylation variation between sexes remains unclear, although it is accepted that females undergo a slower aging rate, as they have a slower methylomic aging rate (Hannum *et al*. 2013) and a later age-related decline (Gur and Gur 2002). In our study, significant differences in global methylation levels between males and females were observed in all CpG types, with hypermethylation at CGI and hypomethylation at CpG shore, shelf, and open sea in females compared to males (Supplementary file 1, Table S11-S14). As we found that the average methylation at CGI increased with age but the average methylation at CpG shore, shelf, and open sea decreased in elders, differences in methylation pattern between males and females may result from a slower methylomic aging rate in females. These results suggest that regions located near to and far from CGIs tend to be methylated and demethylated, respectively, over aging in a sex-specific manner, and that the global sex-specific methylation level may change across the genome.

### Functional annotation of aDMPs

Age-related diseases such as metabolic syndrome, obesity, type 2 diabetes mellitus, and cardiovascular diseases are increasing due to the growing aging population observed worldwide. There are several mechanisms that play important roles in the development of age-related diseases, including epigenetic processes (Franceschi *et al*. 2018). To assess the functional relationship between epigenetics and age-related diseases, genes associated with aDMPs were analyzed by disease association analysis. We investigated 10,596 genes that were determined to have aDMPs for their various disease associations. From these, a total of 6,740 genes have been confirmed to be associated with a disease based on GAD annotation. We identified 22 diseases to be significantly associated with the aDMP genes (FDR < 0.05). The most significant related disorder was tobacco use disorder, with 2,177 related genes (Supplementary file 1, Table S15). It has been shown that smoking can affect DNA methylation, and some of these genes are involved in the risk of age-related diseases such as cardiovascular diseases (Breitling *et al*. 2012; Dogan *et al*. 2014). Our results indicate that the accumulation of epigenetic changes with age were comparable to smoking-associated DNA methylation changes, which imply that smoking can likely change the dynamic of the DNA methylation pattern similarly to the aging process. The GAD analysis revealed other age-related diseases associated with aDMP genes such as type 2 diabetes (Rönn *et al*. 2008), erythrocyte count (Detraglia *et al*. 1974), body mass index and related disorders (e.g., waist circumference and body weight) (Hochberg *et al*. 1995; Poehlman *et al*. 1995), cholesterol (Kreisberg and Kasim 1987), triglycerides (Tucker and Turcotte 2003), cardiovascular diseases (e.g., heart rate, heart failure, stroke, coronary artery disease, blood pressure, and arteries) (Taddei 2009; Breitling *et al*. 2012) and Parkinson disease (Levy 2007) (Supplementary file 1, Table S15). Our results also indicated that age-related DNA methylation modifications in 312 genes were related to alcoholism disorders.

A total of 3,328 genes were found to be associated with KEGG Pathways that have been previously related to aging (FDR < 0.05) (Supplementary file 1, Table S16). For instance, the most significant KEGG pathway was the focal adhesion. The aging process in humans is associated with reduced flexibility of joints and tissue elasticity that is largely caused by alterations in focal adhesion formation on the surface of cells (Arnesen and Lawson 2006). The second significant KEGG pathway was the regulation of the actin cytoskeleton, which plays a fundamental role in cellular pathways (Mooren *et al*. 2012). A total of 142 genes were significantly associated with this KEGG pathway. It has been shown that aging can reduce cytoskeletal integrity, and that overexpression of the heat-shock transcription factor, HSF-1, can function in the preservation of the cytoskeleton (Baird *et al*. 2014). Thus, these results provide evidence that age-associated epigenetic landscape alterations may contribute to disease susceptibility through biological pathways.

### Methylome aging rate and sex-specific methylomic aging rate (AMAR)

To identify sex-specific methylomic aging rates, we compared three different models: i) no differences between males and females (Model 1), ii) differences exist between males and females (Model 2), and iii) there are interactions between sex and age (Model 3), for AMAR over aging in RKHS and BRR models without sex variable. AIC e showed that sex significantly affected the methylomic aging rate in BRR and RKHS (Supplementary file 1, Table S17). As shown in Figure 5, the AMAR was significantly reduced in females compared to males by age (P < 0.01). This result is in agreement with a previous study, which showed a faster aging rate in males (Hannum *et al*. 2013).

**Figure 5.**
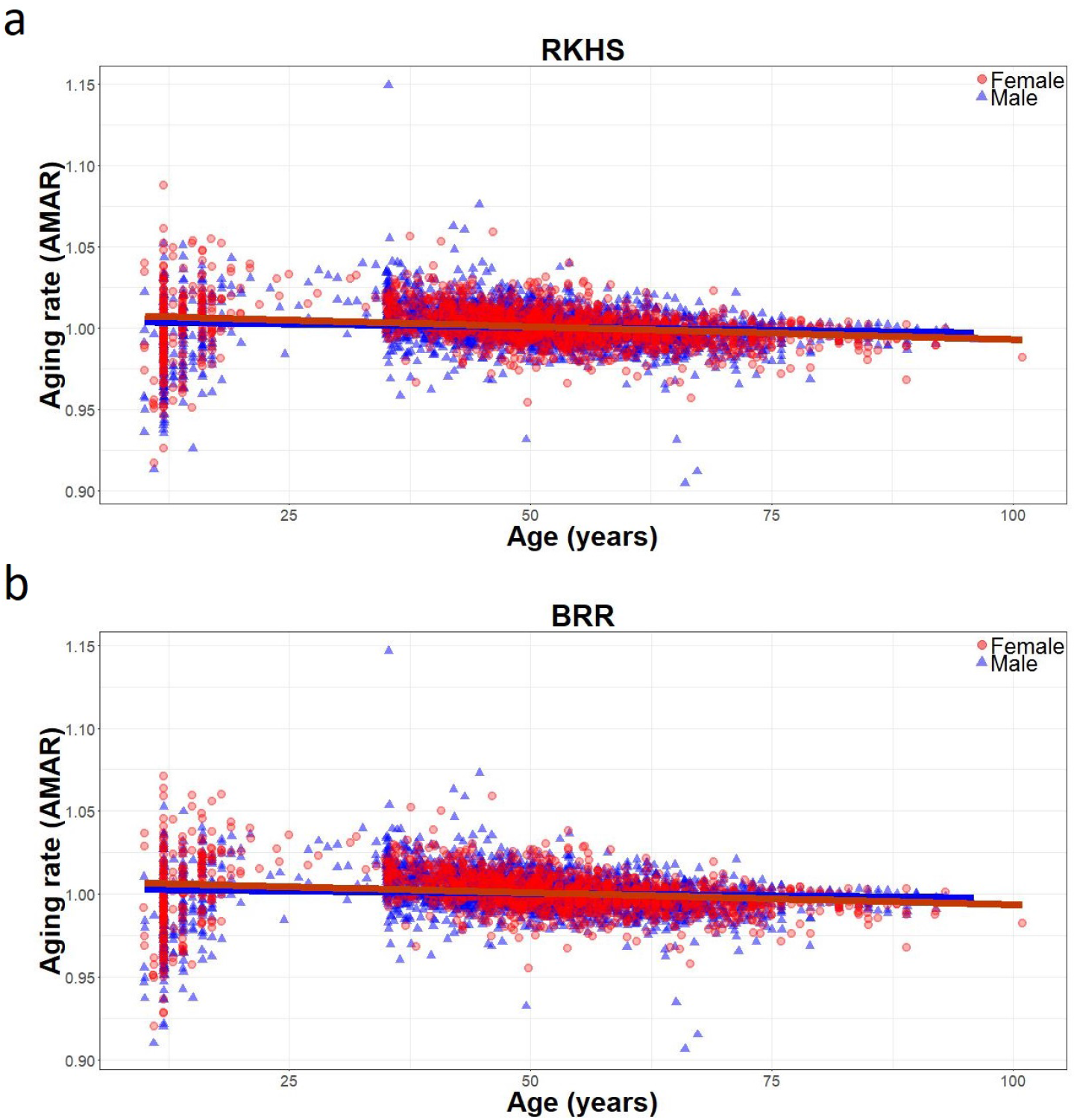
Regression of apparent methylomic aging rate (AMAR) on age in males (blue) and females (red). Models were (a) reproducing kernel Hilbert space (RKHS) and (b) Bayesian Ridge Regression (BRR); both models showed sex-specific AMAR patterns. In both approaches, females had significantly slower AMAR than males (p < 0.01; see Supplementary file 1, Table S17).

In summary, we identified 1,549 suggestive sites, which may influence AMAR producing sex-specific methylation patterns (Supplementary file 2). Genetic and environmental factors can hasten or delay the aging process. Several aging phenotypes have been associated with genetic variation within the FOXO3A gene (Willcox *et al*. 2008), and some of these genetic variants are associated with a slowing or delay in age-related disease (Kenyon 2010). Lifestyle choices such as smoking and physical fitness are among the environmental factors that can influence the aging process (Blair *et al*. 1989). Sex can also affect human longevity, as mortality rates are higher for males, and females tend to live longer than males in most countries (Austad 2006). These observations have led to the search for molecular markers of aging rate, and a faster methylomic aging rate in males than in females has been reported (Hannum *et al*. 2013). Although it is apparent that females can live longer than males, there is a limited number of studies attempting to understand the mechanisms that drive changes in the aging rate in a sex-specific manner. Methylomic assessment of sex-specific age-related methylation sites allows us to understand differences of methylomic aging between sexes. We identified over 1,500 putative methylation sites that could affect variation in methylomic aging rate. This large number of sex-specific methylated aDMPs highlight the complexity of methylomic aging rate.

## CONCLUSIONS

We showed that adjusting methylation beta values for factors, such as spatially cell heterogeneity, can conceal association signals. Our WMP analysis indicated detrimental effects on prediction accuracy from the use of adjusted methylation measurements. Using a small number of methylation sites based on previous studies, by contrast, showed better results when adjusted methylation was used in the prediction model. Here, the use of cell-type corrections should be considered carefully in chronological age prediction. Further research using other traits is important to evaluate cell-type adjustment effects on prediction accuracy. When we considered age-associated methylation sites according to the position in CpG type, a distinct global methylation pattern emerged. This pattern generally showed hypermethylation in CGIs and hypomethylation in off CGIs.

Although BRR and RKHS methods can deal with the problem caused by a number of predictors larger than the sample size, we have shown that choosing age-associated methylation sites according to EWAS for pre-selection of predictors assists in improving prediction ability. The exclusion of not-associated markers resulted in a better prediction model. Using a subset of methylation sites without heteroscedasticity, a widespread practice, can drastically reduce the number of predictors while producing a reasonable prediction accuracy. Further research on preselection methods may help to find even more accurate prediction models. Our results illustrated that age prediction based on RKHS and BRR using all or EWAS/BPtest subset methylation sites is very effective relative to using a small number of candidate sites based on previous studies.

We found that methylation sites can display a sex-specific methylation rate. These sites may help explain the increased methylomic aging rate for males. We found sex-specific methylomic aging processes, but much remains to be learned in understanding the molecular mechanisms underlying sex-specific aging processes.

## Availability of data and materials

The Illumina 450K methylation array datasets analyzed during the current study are publicly available in GEO with accession numbers “GSE55763”, “GSE40279”, “GSE50660” and “GSE56105”.

## Authors’ contributions

MAR and MM conceived and planned the experiments. MAR carried out the experiments with support from FBL, SSA and SFM. MAR and SFM wrote the manuscript. HK and DG contributed to the experimental design of the study and participated in the manuscript writing. All authors read and approved the final draft of the manuscript.

## Ethical approval and consent to participate

We confirm that all methods were performed in accordance with the relevant guidelines and regulations.

## Competing interests

The authors declare no competing interests.

